# A simple and robust procedure for preparing graphene-oxide cryo-EM grids

**DOI:** 10.1101/290197

**Authors:** Eugene Palovcak, Feng Wang, Shawn Q. Zheng, Zanlin Yu, Sam Li, David Bulkley, David A. Agard, Yifan Cheng

## Abstract

Graphene oxide (GO) sheets have been used successfully as a supporting substrate film in several recent cryogenic electron-microscopy (cryo-EM) studies of challenging biological macromolecules. However, difficulties in preparing GO-covered holey carbon EM grids have limited its widespread use. Here, we report a simple and robust method for covering holey carbon EM grids with GO sheets and demonstrate that these grids are suitable for high-resolution single particle cryo-EM. GO substrates adhere macromolecules, allowing cryo-EM grid preparation with lower specimen concentrations and providing partial protection from the air-water interface. Additionally, the signal from images of the GO lattice beneath the frozen-hydrated specimen can be discerned in many motion-corrected micrographs, providing a high-resolution fiducial for evaluating beam-induced motion correction.

## Introduction

Recent technological breakthroughs have made single particle cryogenic electron microscopy (cryo-EM) a versatile and routine method for structure determination of macromolecules at high-resolution (Bai et al., 2015; Cheng, 2015). With automated data acquisition (Mastronarde, 2005; Suloway et al., 2005) enabled by stable high-end electron microscopes equipped with direct electron detection cameras and streamlined image processing software (Kimanius et al., 2016; Punjani et al., 2017), structure determination by single particle cryo-EM has never been easier. What remains more or less unchanged is the plunge freezing technique (Dubochet et al., 1988), which works well for many samples, particularly structurally stable ones. For some fragile complexes, however, preparing good frozen-hydrated cryo-EM grids with intact and monodispersed particles is often a challenging task and a main bottleneck to cryo-EM structure analysis. It has been suggested that exposing protein samples to an air-water interface during plunge freezing can damage fragile protein complexes or induce preferred orientations in thin vitreous ice (Glaeser, 2018; Glaeser and Han, 2017). Additionally, some samples may prefer to stick to the carbon matrix of holey grids instead of being suspended in the vitreous ice spanned by the holes (Snijder et al., 2017; Zhao et al., 2015b). A common approach to mitigate these problems is to apply a continuous thin layer of substrate to the holey carbon grid, evening the distribution of particles inside the holes and holding protein samples away from the air-water interface (Glaeser, 2018; Han et al., 2016; Russo and Passmore, 2014; Williams and Glaeser, 1972).

Amorphous carbon is the most commonly used substrate (Grassucci et al., 2007), but it adds significant background noise to particle images, limiting its use to relatively large particles such as ribosomes (Gao et al., 2007). Other substrates include monolayer sheets of graphene (Pantelic et al., 2011; Russo and Passmore, 2014) and two-dimensional crystals of streptavidin (Han et al., 2017; Wang et al., 2008). More recently, monolayer sheets of graphene oxide (GO) were introduced as a substrate (Bokori-Brown et al., 2016; Boland et al., 2017; Pantelic et al., 2010). Compared to other options, GO sheets are nearly electron transparent, hydrophilic enough to adhere macromolecules from dilute solutions, inexpensive to purchase or synthesize, and amenable to functionalization (Chen et al., 2012; Pantelic et al., 2010). However, obtaining EM grids evenly covered with one or several layers of GO sheets has not been easy. In our experience, our attempts to use the previously reported drop-casting method (Pantelic et al., 2010) mostly produced grids with irregular coverage of GO sheets over the holes. Only a small percentage of holes were covered by one or a few layers of GO sheets, with the majority of the grids either covered with multi-sheet aggregates or lacked GO entirely. Because we regularly need to screen tens of cryo-EM grids before finding conditions suitable for automated data acquisition and high-resolution structure determination, reproducibility and ease of manufacture are key considerations for any substrate.

To improve the usable area on GO covered grids, we established a simple and robust surface assembly procedure for evenly covering holey carbon EM grids with one to very few layers of GO sheets. These GO grids are suitable for high-resolution single particle cryo-EM studies of biological macromolecules. We prepared frozen-hydrated archaeal 20S proteasomes using such GO-covered Quantifoil EM grids and collect a dataset resulting in a 2.5Å resolution 3D reconstruction, comparable to our best previous results in freestanding vitreous ice (Li et al., 2013; Zheng et al., 2017). We also confirmed that particles are concentrated close to the GO sheet, away from the air-water interface. In addition, lattice images of the graphene oxide film recorded together with frozen-hydrated 20S proteasome particles provide information for evaluating beam-induced motion correction.

### Fabricating GO-covered holey carbon cryo-EM grids

Ideally, a GO covered grid would be completely covered in a single monolayer GO sheet without any wrinkled regions or GO aggregates. Direct application of an aqueous suspension of GO sheets to a glow-discharged EM grid (drop-casting) (Pantelic et al., 2010) tends to leave many regions of the grid uncovered and deposits high-contrast multi-sheet aggregates over many others. Initially we speculated that our commercial GO suspension might have deteriorated with age, so we used bath sonication to break up weakly aggregated sheets followed by centrifugation to isolate mostly large single GO sheets. This treatment reduces the presence of multi-sheet aggregates, but does not greatly improve coverage uniformity.

GO sheets are sufficiently hydrophobic to be enriched at air-water interfaces (Kim et al., 2010). We used this property to assemble a thin, mostly continuous film of GO sheets (typically one to three layers) at the surface of a dish of pure water. By draining the water, the assembled GO film is then slowly lowered onto submerged holey carbon EM grids with their holey carbon film sides facing up (Figure 1A). In agreement with previous reports (Cote et al., 2011; Kim et al., 2010), we found that methanol-dispersed GO sheets spread and float easily on a pure water subphase. Once completely dried, GO sheets stably adhere to holey carbon films and remain adhered during blotting. The boundaries of individual GO sheets on EM grids can be directly observed in the transmission electron microscope (TEM) (Figure 1B). Because each individual sheet produces a hexagonal pattern of Bragg peaks, the number of GO sheets spanning any given hole can be discerned in diffraction mode (Figure 1C). Such a diffraction pattern with sharp high-order spots indicates a long-range periodicity of the GO lattice over the hole. While GO grids can acquire surface contaminants which render them less hydrophilic, they can be cleaned without damage by brief glow-discharge in air (five to ten seconds) immediately before use. A detailed protocol for fabricating GO grids by surface assembly is provided in the supplementary methods.

**Figure 1.**
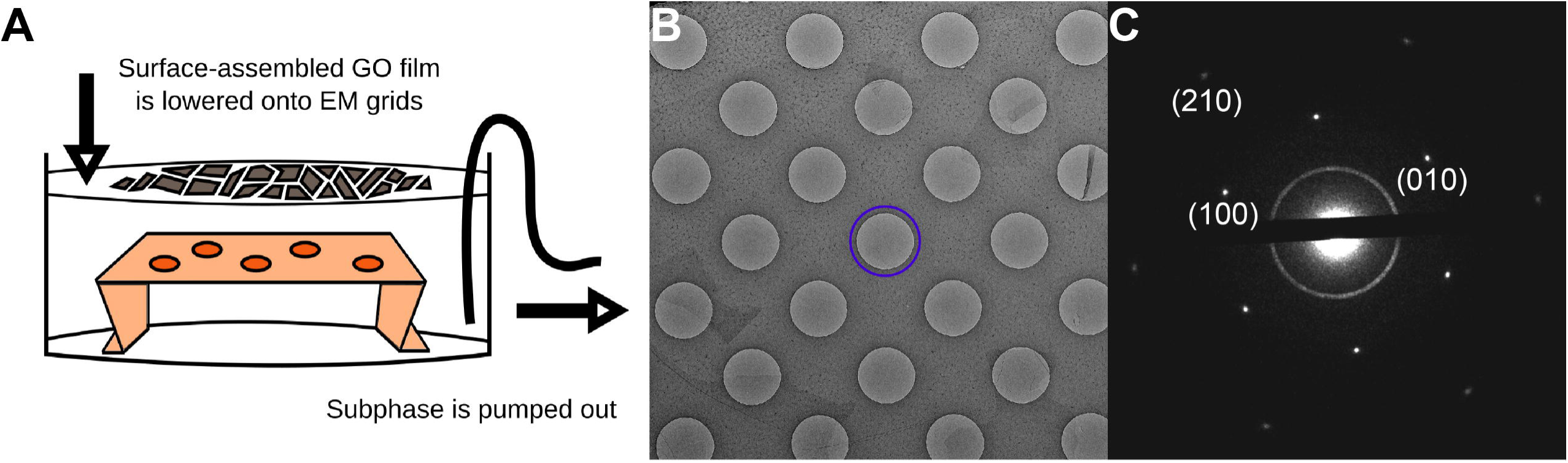
EM grids can be evenly coated with GO sheets using surface assembly. **A**. Schematic of the apparatus used for surface assembly of thin GO films and subsequent deposition onto holey carbon EM grids. **B**. Low magnification image of a holey carbon (Quantifoil) grid covered with a thin film of GO sheets. A hole covered by a single GO sheet is circled in blue. **C**. Electron diffraction pattern of the hole in B. Without a scrolled edge, it is hard to discern by image contrast alone whether a hole has a GO sheet spanning it. Instead, the diffraction pattern shows unambiguously the hexagonal pseudo-crystalline lattice of each GO sheet.

Suspensions of GO sheets can vary significantly in the lateral sheet size and degree of oxidation. We obtained the best results with home-made GO suspensions optimized for large sheet size, as these sheets show fewer high contrast ‘edges’ in images (Marcano et al., 2010). Even so, surface assembly is robust to these variations and works well with both commercially-available GO suspensions and home-made GO sheets. The only important parameter to optimize with our protocol is the amount of GO applied to the surface. We typically make one or two test grids and ensure satisfactory coverage by screening in a transmission electron microscope before producing a large batch of grids.

### Single particle cryo-EM of archaeal 20S proteasome on a GO grid

Using the archaeal 20S proteasome as a test specimen, we evaluated the practicality of using GO grids for high-resolution single particle cryo-EM. As previously reported, the commonly used plunge freezing procedure works well for GO grids, though we have found that longer blotting times (10-30 seconds) are often preferred. For the 20S proteasome, achieving an optimal particle distribution required a specimen concentration approximately ten times lower with GO (0.05 mg/mL) than without (0.5 mg/mL) (Figure 2A, Supplementary Figure 1A). We confirmed that nearly all 20S proteasome particles physically adhere to the GO surface using cryogenic electron tomography (Figure 2B, Supplementary Figure 1C-D). Based on the locations of the few proteasomes and gold nanoparticle fiducials not adhered to the GO face, we estimate the thickness of the vitreous ice in this hole to be about 65 nm, while the deviation in the Z-location of 3D template-matched 20S proteasome particles is only about 5 nm. This suggests that 20S proteasomes on GO are protected from the air-water interface and lie on a common plane, a clear benefit when estimating micrograph defocus.

**Figure 2.**
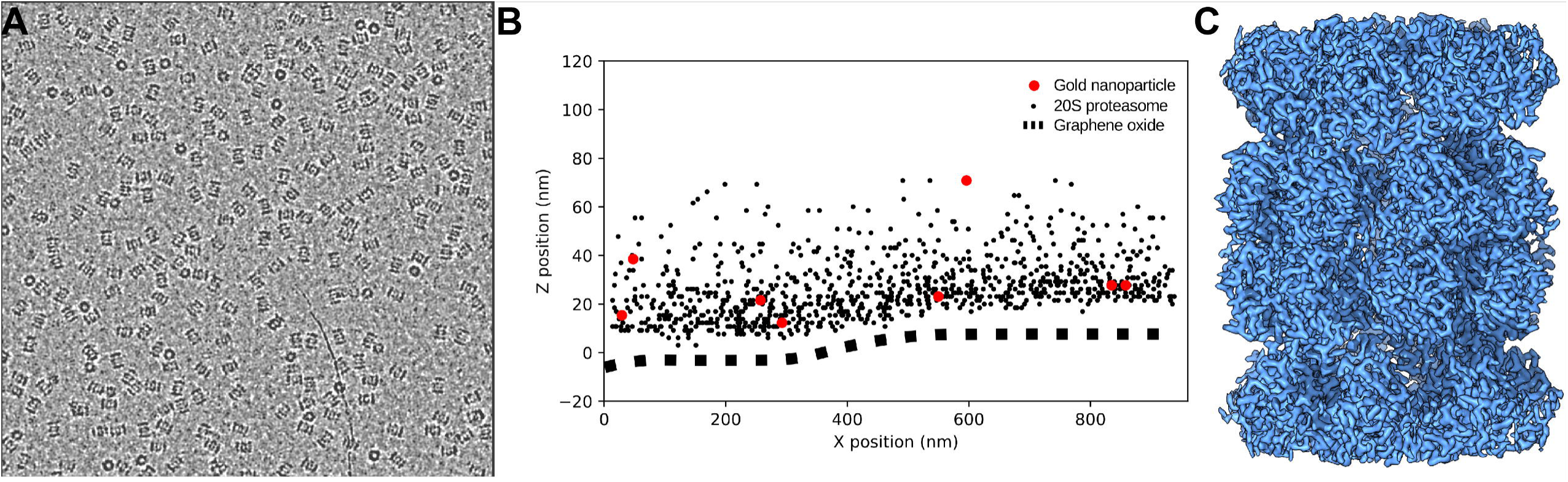
High-resolution single-particle cryo-EM on GO grids. **A**. Micrograph of 20S proteasome particles over a GO support film. The GO film adds minimal background contrast. The specimen was applied at a concentration 0.05mg/mL, approximately ten times less than we typically use for this specimen. **B**. 3D localization of 20S particles from cryogenic electron tomography (cryo-ET). This tomogram was taken from a different GO grid than the one we used for the single-particle results: the 20S proteasome concentration here was higher and the GO substrate was far thicker (5-8 sheets). These conditions enabled us to identify how densely the GO surface could be coated with macromolecular specimen and to discern the position of the GO layer within low-resolution tomogram. Fiducial markers and the location of the GO sheets were picked manually, while 20S proteasome particles were located in 3D using template matching. The GO is not flat but ripples and bends. The vast majority of 20S proteasomes apparently adhere directly to the GO sheet. All tomographic processing was performed using the IMOD software suite (Mastronarde, 2005). **C**. 3D reconstruction of the 20S proteasome on a GO support resolved at 2.5Å.

To compute a 3D reconstruction by single-particle analysis, we used the SerialEM automated data acquisition procedure to collect 740 dose-fractionated stacks. When choosing areas for data acquisition, we did not attempt to distinguish if holes were covered by GO or not. In this dataset, approximately 73% of the collected images were of high-quality, judged from the particle distribution and lack of numerous high-contrast GO edges. 10% of micrographs have poor ice quality likely unrelated to the GO, 10% of the holes are missing the GO support, and 7% of the holes had suboptimal coverage (usually too many overlapping sheets). Beam-induced motion was corrected using MotionCor2 (Zheng et al., 2017). We did not notice any significant differences in the magnitude of beam-induced motion when comparing images of holes with no GO, with thin optimal GO, or with thick suboptimal GO.

For 3D reconstruction, we included only particles collected from holes covered with GO sheets and did not attempt to computationally remove the periodic graphene lattice, as might be done when using other crystalline lattice supports such as streptavidin crystal grids (Han et al., 2016; Wang et al., 2008). From this dataset, we determined a 3D reconstruction of the 20S proteasome at a resolution of 2.5Å (Figure 2C, Supplementary Figure 1E). The density map is visually indistinguishable in quality from our previously reported map of the 20S proteasome in freestanding vitreous ice but used fewer particles in the final reconstruction (117,578 particles on GO vs. 187,011 in freestanding vitreous ice (Zheng et al., 2017)). This suggests that despite the added background contrast and the micrographs lost due to defects in the GO grid fabrication, GO poses no significant barrier to achieving high-resolution reconstructions.

### Evaluating correction of beam-induced motion

GO sheet typically maintains a long-range periodicity (Figure 1C) that can tolerate the amount of electron beam radiation used to image the biological sample (Supplementary Figure 2). For a typical image recorded as a movie stack of subframes, only weak spots are seen in the sum image before motion correction (Figure 3A). These peaks can also be observed in short three-frame averages during the exposure, suggesting that the GO lattice is not significantly deteriorated by radiation damage but is primarily blurred by beam-induced motion (Supplementary Figure 2). After full motion correction with MotionCor2 (Zheng et al., 2017), the power spectrum show one complete set of hexagonally arranged peaks around 2.1Å resolution, beyond the physical Nyquist frequency of the K2 camera (2.43Å at the selected magnification) (Figure 3B). These peaks signal the first order reflections of the underlying GO lattice, with each hexagonal set of peaks corresponding to a single GO sheet. Reflections are typically very sharp, with an average half width of 10 Fourier pixels, suggesting that the vitrified GO lattice covering the image area maintains a long-range periodicity.

**Figure 3.**
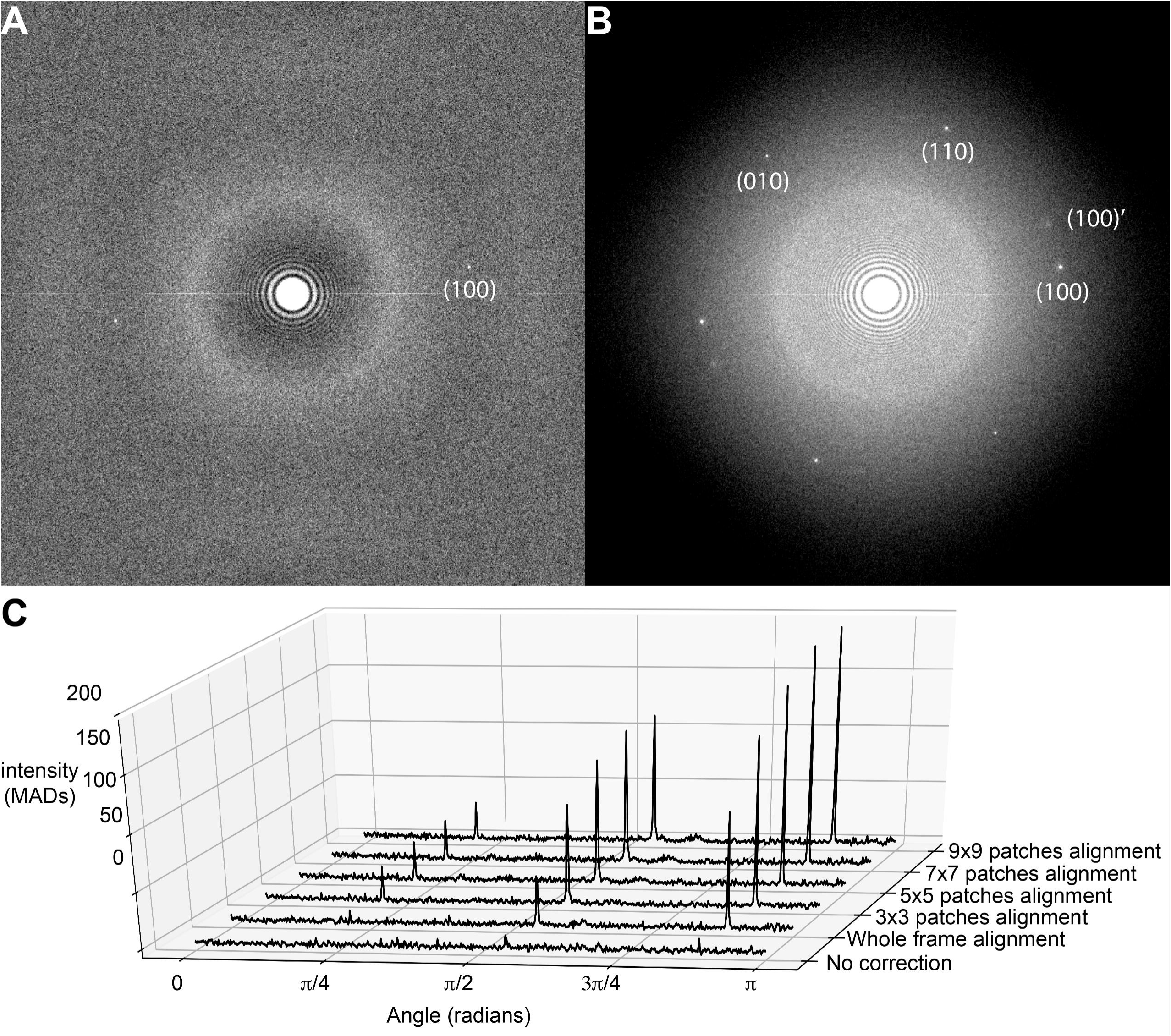
Evaluation of motion correction using GO peak intensities. Power spectra from the image in Figure 2A before (**A**.) and after (**B**.) motion correction. Power spectra were calculated by averaging the periodograms of overlapping 512×512 pixel windows using a script in Python. **C**. GO peak height increases as the number of local motion patch trajectories calculated by MotionCor2 is increased. To generate 1D radial profiles showing the GO peaks with respect to the background, the aforementioned power spectrum was transformed into polar coordinates by cubic spline interpolation and a radial band of intensities from 2.0Å to 2.2Å was extracted. The maximum intensity component in this band was taken for each angle sampled in the half-circle 0 to π. This was done because the actual spatial frequency of the GO peak deviated from the expected position at 2.13Å, likely due to residual specimen tilt or uncorrected anisotropy in the magnification system (Zhao et al., 2015a). To compare 1D profiles between different motion correction schemes, the intensities in each 1D profile were scaled according to the median absolute deviation (MAD), which is a robust measure of scale. All calculations were performed in python with the scientific computing library scipy, with code available on request.

Assuming the primary source of deformation in the GO lattice image is beam-induced motion, the intensity and sharpness of these recovered peaks should correlate directly with the quality of motion correction, and thus can be used to directly evaluate the quality of the motion correction performed on a dose-fractionated image stack. We tested this by comparing GO peak heights before and after using MotionCor2 (Zheng et al., 2017) to correct global as well as local motions.

In MotionCor2 (Zheng et al., 2017), global motions are corrected by iteratively refining translational shifts for each frame, while nonuniform local motions are corrected by fitting numerous trajectories of local motions to a time-variant polynomial function. This polynomial estimates the instantaneous shift for any location in the image at any time point in the exposure, allowing local motions to be corrected smoothly at every image pixel. The number of image patches where local motion trajectories are measured is a free parameter in MotionCor2: a coarse grid of patches (i.e. 3×3 patches) will provide fewer measurements, but they should have high signal-to-noise ratio (SNR), while a finer grid (9×9 patches) will provide a finer measurement of localized motion, but at the expense of increased computation and a lower per-patch SNR.

GO peaks are nearly absent in the power spectra before motion correction, but they are readily visualized after global motion correction (Figure 3A and C). Applying local motion correction, we observe a trend where increasing the number of patches significantly increases GO peak intensity. For this particular image, using 7×7 patches increased the peak intensities approximately 200% with respect to the global alignment, while 9×9 patches provided no additional significant increase. While the intensities of all three distinct GO lattice peaks increase with better motion correction, the amount of improvement is different for each. We speculate that this could occur if the image still contains uncorrected motion, such as that occurring within each frame and if this motion were orthogonal to the direction of the foreshortened peaks.

## Conclusions

We have established a simple and robust procedure for covering holey carbon EM grids with thin films of GO sheets and have demonstrated the utility of the resultant grids for determining 3D reconstructions of macromolecules at high-resolution. We also showed that GO peaks in the image power spectrum are sensitive fiducial markers for the quality of the beam-induced motion correction. By simplifying their production, we anticipate that our method will make the benefits of GO grids more widely accessible.

Nevertheless, we believe the GO grids described here are only one step towards a more universal substrate for high-resolution single-particle cryo-EM. In the course of testing this protocol, we have attempted to use GO grids in several challenging, on-going cryo-EM projects. In some cases, macromolecular particles were bound and concentrated onto the GO surface in a native state like we report here for the 20S proteasome. In other cases, particles were not visibly bound to the GO or appeared denatured. This could occur if the proteins have no affinity for GO, as they would then have no protection from the air-water interface. There have also been reports that GO itself can destabilize protein structure and compromise enzymatic activity (Bai et al., 2017). In either case, an obvious solution is to functionalize the GO surface, increasing its affinity for interactions that preserve protein structure while passivating residual non-oxidized hydrophobic domains that may promote denaturation. GO has abundant epoxide groups on its surface that are amenable to such functionalization. We are actively working along these lines to improve the applicability of GO grids to fragile specimens.

Even if proteins interact favorably with GO and are protected from the air-water interface, they can still denature when directly exposed to filter paper during blotting. To protect GO-bound proteins from possible deleterious filter paper interactions, we now routinely apply sample to the back side of the grid. During blotting, the grid bars should help prevent the filter paper from directly contacting the GO-bound proteins (Supplementary Figure 3). This method was originally used by electron crystallographers who called it ‘back-side injection’ (Gyobu et al., 2004). We have found that long blotting times (20-30s) are needed to get appropriately thin vitreous ice with back side application.

If a protein specimen is properly adhered to GO grids, we expect that imaging conditions should not be worse than if no substrate is used, and may be better. Particles bound to a common surface should have locally-correlated heights, improving the accuracy of defocus estimation. We noticed that the GO peaks in the power spectrum do not follow precise hexagonal symmetry, and that the lattice spacing for each of the three peaks vary slightly around 2.1Å. This can be caused either by anisotropic magnification of the microscope or a slight tilt of the specimen, or a combination of the both. If the anisotropic magnification of the microscope is precisely calibrated, it is possible to derive the tilt angle and orientation of the specimen, facilitating a better local defocus determination. At a resolution of ∼2.5Å, we did not notice any influence of the underlying GO lattice on particle alignment or classification for the 20S proteasome reconstruction. We therefore did not attempt to computationally remove the underlying GO lattice. It may become necessary to do so if the resolution of a reconstruction is beyond the GO lattice space, ∼2.1 Å.

Finally, we showed that subtle improvements in beam-induced motion correction can be detected by examining the intensity of GO peaks present in the power spectra of differently motion corrected micrographs. This result suggests that directly using the intensity of the GO peaks as an objective function for beam-induced motion correction might give improved results. The principal challenge here is that lateral shifts larger than the unit cell of the GO lattice (2.13Å) will give degenerate solutions. Fortunately, after rigid body motion is corrected, residual local motions tend to be only a few angstroms. If used to locally refine the trajectory estimated from the lower-resolution image features, we expect the GO peaks could improve the accuracy of local beam-induced motion correction, recovering the highest-resolution features captured in dose-fractionated electron micrographs.

## Acknowledgement

This work is supported in part by NIH grants R01GM082893, R01GM098672, R01HL134183, P50GM082250, P01GM111126, and 1S10OD020054 to Y.C. and NIH U54 CA209891 and UCSF Program for Breakthrough Biomedical Research to D.A.A. D.A.A and Y.C. are Investigators of Howard Hughes Medical Institute.

---

**Supplemental Figure 1:**
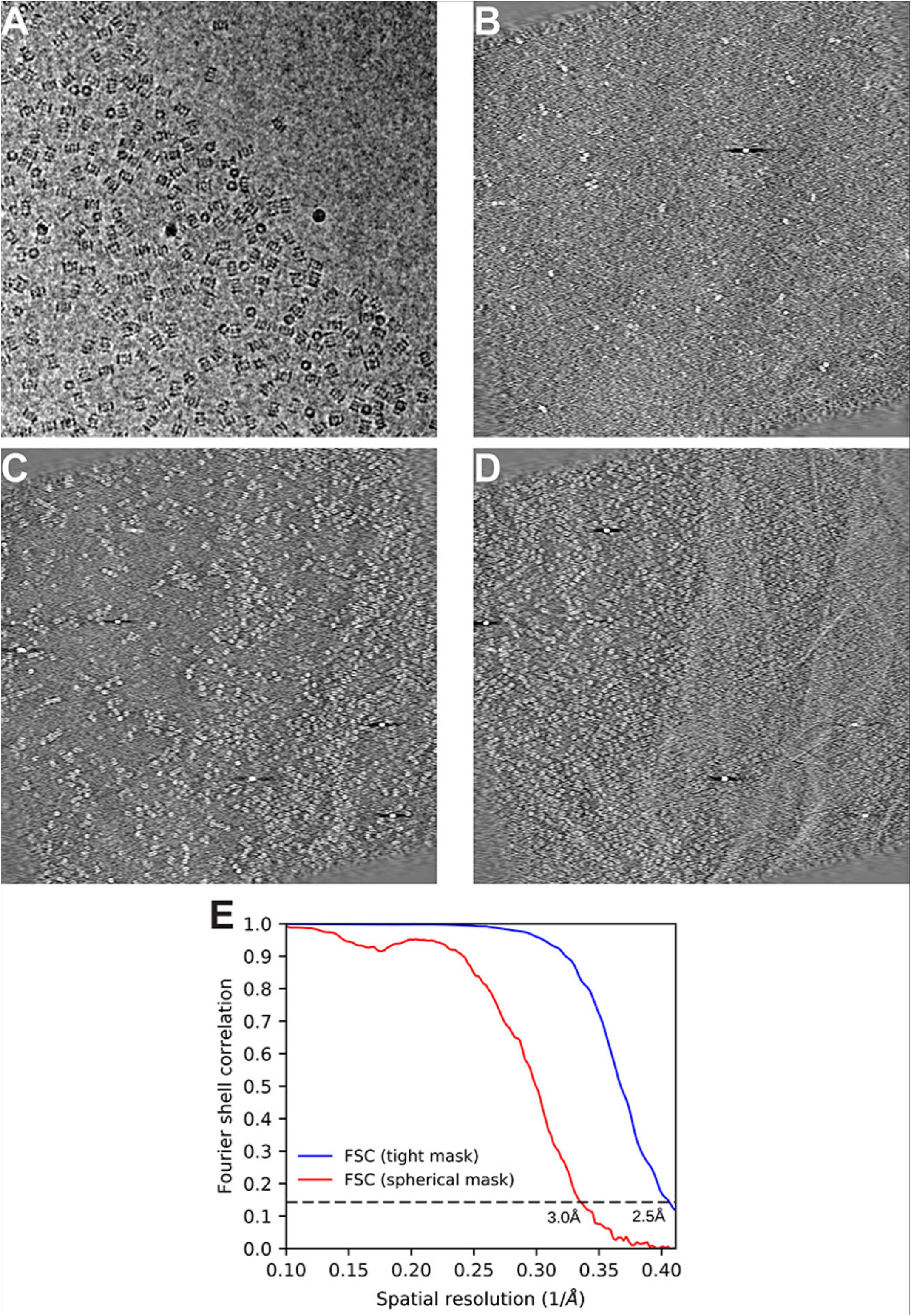
Cryo-EM of proteasome particles prepared using GO covered grid. **A**. Image of a hole partially covered with a GO. Fewer proteasomes are observed in the upper right-hand corner where the GO support is absent. The additional background contrast added by the GO substrate is barely perceptible. **B-D**. Projections through Z-slices of the tomogram summarized in Figure 2B. The slices depict **B**. the air-water interface (58-80 nm away from the lowest apparent point of the GO substrate), **C**. the higher part of the GO surface where bound proteasomes are observed (23-39nm), and **D**., the lower part of the GO surface where bound proteasomes are observed (8-23nm). **E**. Gold-standard FSC curves for the 20S proteasome, calculated from independently reconstructed half-maps with a spherical mask and with a tight mask.

**Supplemental Figure 2:**
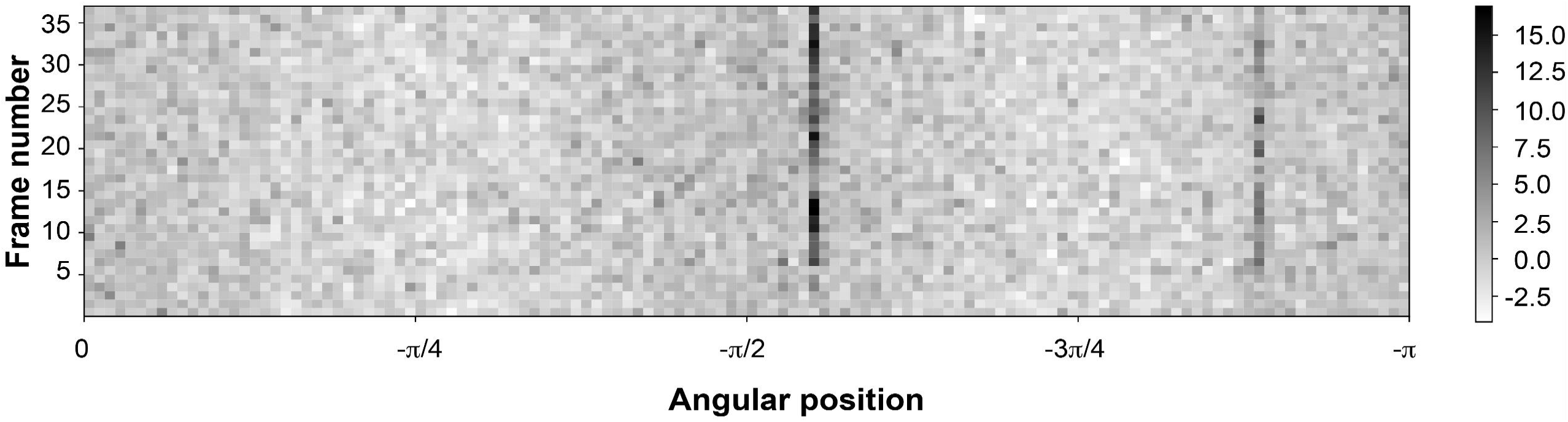
GO peak intensity is not radiation sensitive. From the dose-fractionated stack used in Figure 3, we calculated running three-frame average images without applying motion correction. Each row shows the 1D radial profile for a three-frame average image, calculated identically to those in Figure 3C. Two of three GO peaks are visible in most three-frame average images. While the peaks are not detected at the beginning of the exposure when beam-induced motion is greatest, they are not obviously reduced at the end of the exposure. The color scale represents the MAD.

**Supplemental Figure 3:**
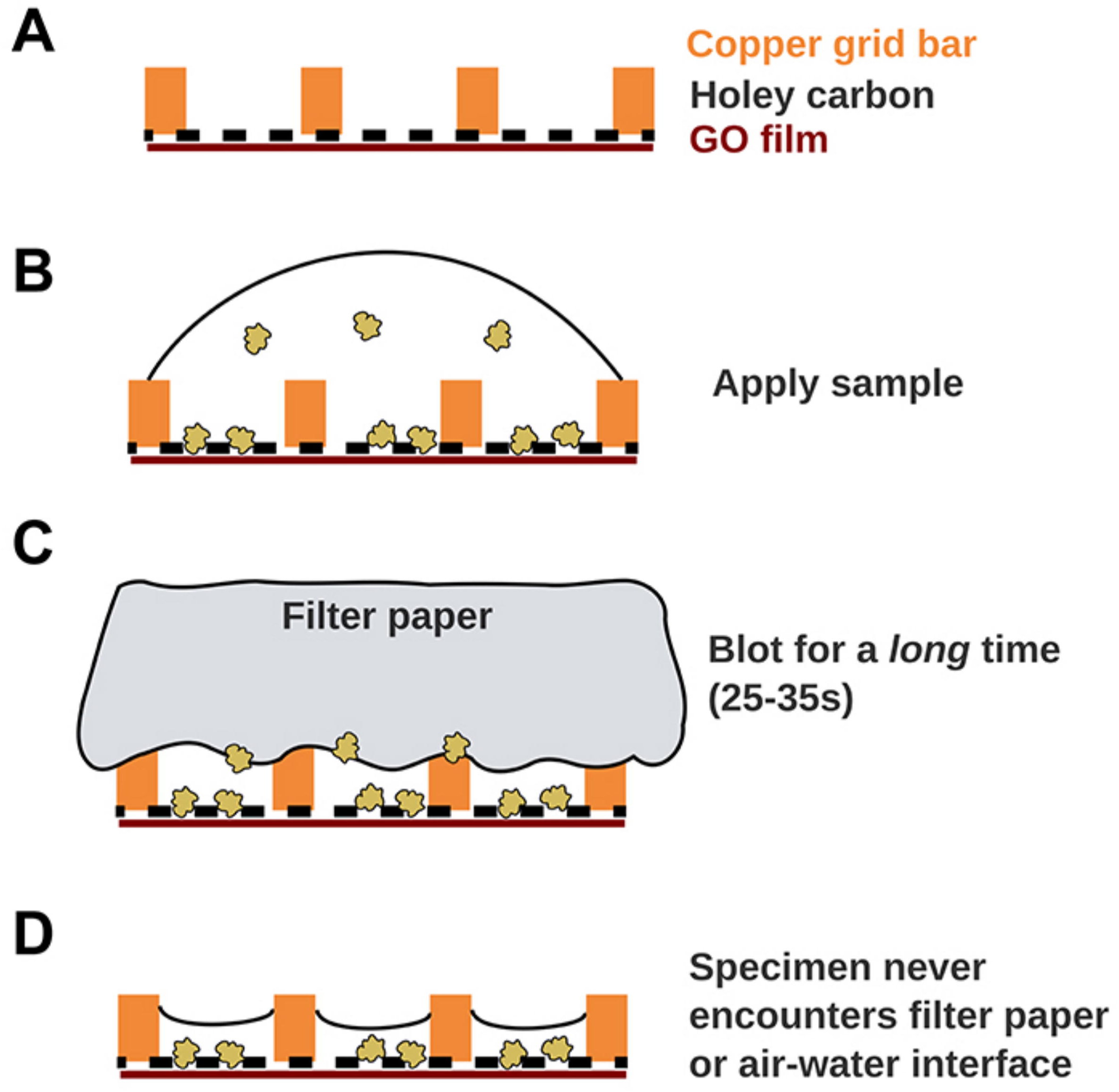
Schematic representation of the back-side application technique.

## Supplementary Materials

### Methods and Materials

Graphene oxide (GO) grids were made on Quantifoil 200 mesh R 1.2/1.3 holey carbon grids according to the protocol described below. The graphene oxide suspension was purchased from Sigma Aldrich (777676 ALDRICH). GO grids were used for cryo-EM once they had completely dried, several hours after they were made.

*Thermoplasma acidophilum* 20S proteasomes were expressed and purified as previously described (Li et al., 2013) and stored in aliquots at −80°C. An aliquot of 20S proteasome was diluted to 0.05 mg/mL in buffer containing 25mM Tris pH=7.5 and 150mM NaCl. 2.5uL of 20S proteasome was applied to a GO grid and allowed to incubate for 30S in the chamber of a Mark III Vitrobot at 100% humidity. The specimen was blotted for 6s with Whatman #1 filter paper and plunged into liquid ethane.

Cryo-EM grids were loaded into a TF30 Polara microscope equipped with a Gatan K2 camera operated in super-resolution mode. Automated data acquisition was performed with SerialEM and 740 micrographs were corrected. Beam-induced motion was corrected in MotionCor2 (Zheng et al., 2017). Per-frame electron dose weighting was performed with MotionCor2 using a nominal electron dose of 1.2 e^−^/A^2^*frame and the radiation damage model derived in (Grant and Grigorieff, 2015). Micrographs were visually inspected and segregated into optimal GO coated (73%), overcoated GO (10%), poor ice (10%), or uncoated (7%) images. 540 micrographs with optimal GO coating were selected for continued processing. CTF parameters were estimated with Gctf (Zhang, 2016) and particles were picked with Gautomatch (https://www.mrc-lmb.cam.ac.uk/kzhang/Gautomatch/) using a template of a 20S proteasome side view. Mispicked particles were removed after 2D classification in cryoSPARC (Punjani et al., 2017)and subjected to homogeneous 3D refinement with D7 symmetry. Masking and FSC estimation were performed automatically in cryoSPARC. The resulting map was compared visually to previous 20S proteasomes maps from our lab with UCSF Chimera.

