## Supplementary Materials for "A simple and robust procedure for preparing graphene-oxide cryo-EM grids"

*Protocol by Eugene Palovcak and Feng Wang - 7 February 2018*

The basic procedure is to form a thin film of GO sheets at the air-water interface of a solution in a glass dish. This solution is termed the 'subphase.' EM grids are submerged in the subphase, sitting on a mesh platform. A peristaltic pump with a line into the dish slowly drains the subphase, lowering the GO film onto the face of the EM grids. When the EM grids dry, the GO tightly adheres to the carbon foil.

While assembling thin films of GO onto air-water interfaces is common in material science, it usually carried out in an expensive Langmuir-Blodgett trough equipped with a surface tensiometer. This allows precise control over the thickness and packing of the GO monolayer film. Since most biochemistry labs do not have ready access to a Langmuir-Blodgett trough, this protocol improvises with a petri dish and a peristaltic pump. The amount of GO applied to the surface must then be optimized in advance, and varies depending on the average lateral size and the degree of oxidation of the stock GO sheets. For this reason, the protocol suggests making several test grids with different amounts of applied GO.

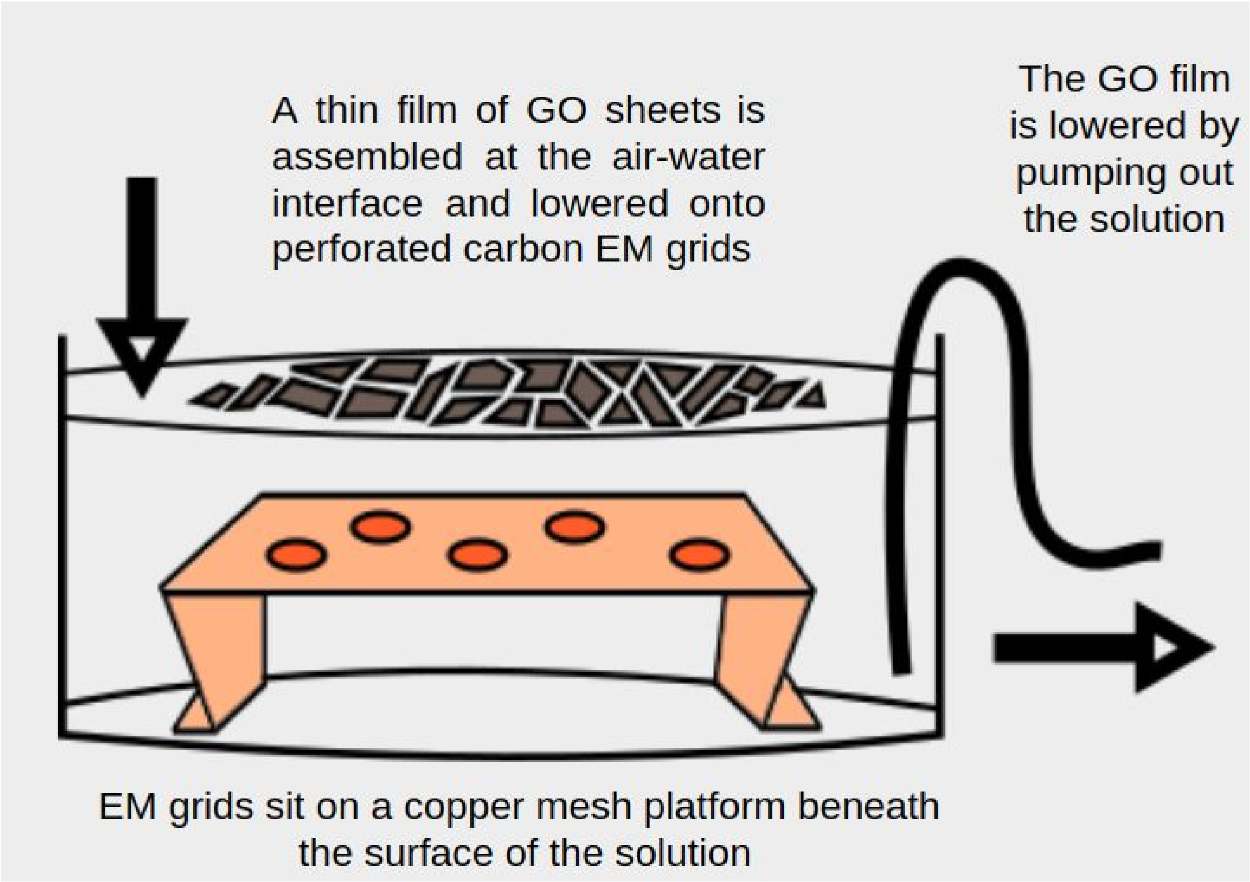

*Materials*
GO dispersed in $H_2O$ (homemade or commercial, such as Sigma 777676 at 4 mg/ml)
Methanol
Quantifoil R1.2/1.3 Cu 200 mesh grids (rinsed in chloroform to remove any plastics)
Chloroform
Ethanol

*Equipment*
Bath sonicator
Spectrophotometer
Glow discharger (Pelco EasiGlow)
Table-top peristaltic pump
6 cm petri dish
Copper mesh (20 mesh), bent into a small 4x4 centimeter platform
Filter paper

*GO solution pre-treatment*
If the stock solution of GO sheets is old, there may be large multi-sheet aggregates that disrupt otherwise even coverage. To remove these aggregates, the GO can be briefly sonicated and then centrifuged to remove the largest, still-aggregated particles. If the GO solution is relatively new, these steps should probably be omitted.

1. Prepare a dispersant solution of 1:5 $H_2O$:Methanol. In a 50 mL conical tube, add 8 mL of $H_2O$ and 40 mL of methanol.
2. Dilute stock solution of GO with dispersant to 0.2 mg/mL to make 10 mL of GO solution. Exact concentration is less important here than the total amount applied. If GO stock solution is homemade or relatively new, proceed to '*Setting up the coating apparatus*'

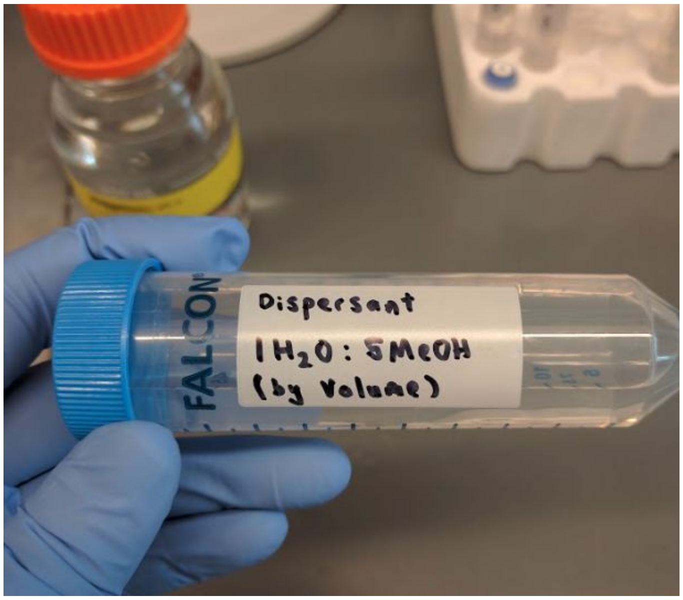


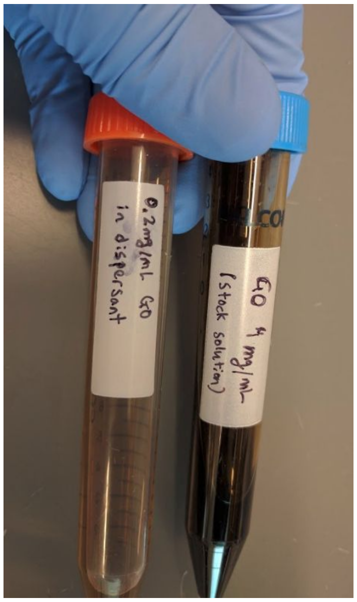

3. Sonicate in a bath sonicator for 10 minutes to break up aggregates of GO sheets.

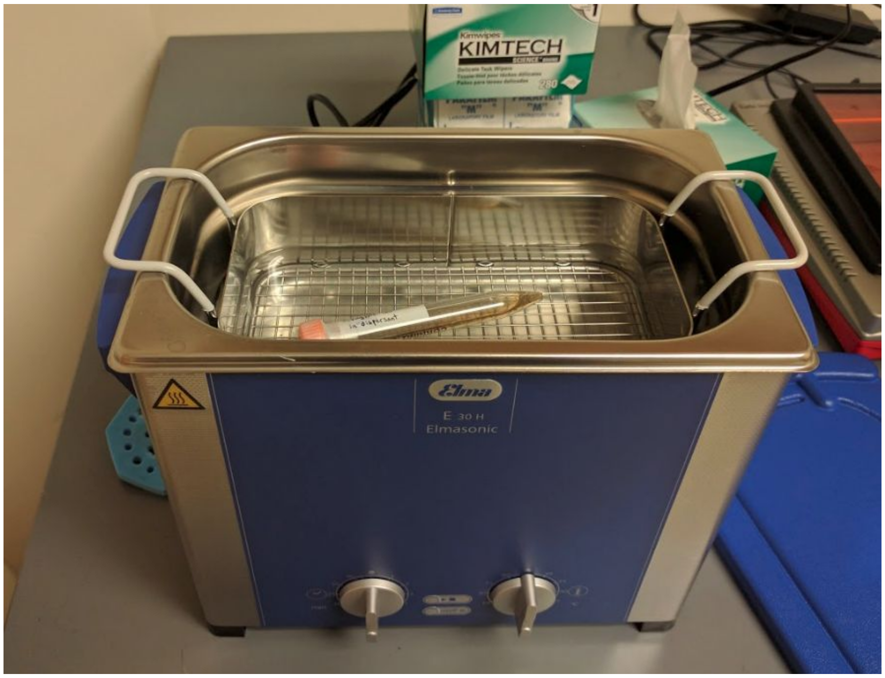


4. Aliquot 9mL into 6x1.5mL eppendorf tubes and centrifuge at 4000xg for 10 minutes.

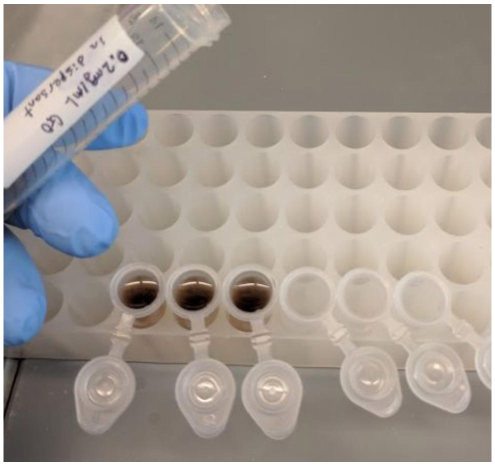


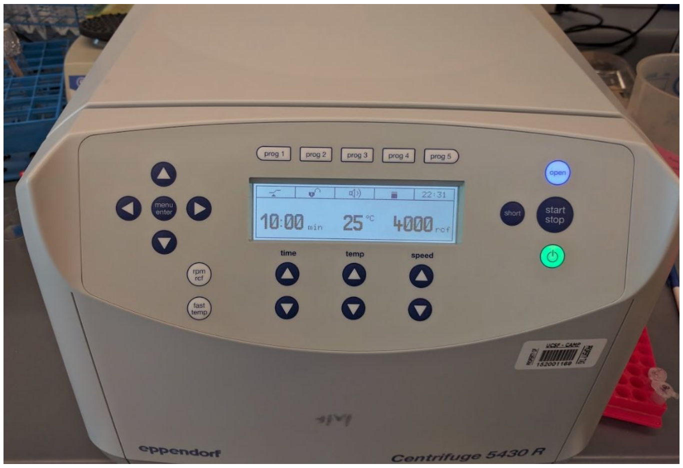


5. Carefully remove the supernatant, which contains the smallest fragments of GO. Resuspend the pellets in 500 uL dispersant. This solution contains the desired large sheets of GO.

6. Sonicate the GO solution for 2 more minutes.

7. Centrifuge for 1 min at 500 xg to pellet residual aggregates and transfer to a new tube.

8. Dilute the GO solution back to 0.2mg/mL with dispersant using a spectrophotometer. The UV-vis spectrum of GO has a shoulder at 305 nm that can be used to measure the concentration by Beer's law. From a 4 mg/ml stock solution, the extinction coefficient at 305 nm is 21.2 ml/mg*cm. Diluting back to 0.2 mg/mL can also be done by visual comparison to a solution diluted from the stock at some expense of reproducibility. The yield is typically about one milliliter of 0.2 mg/mL GO solution.

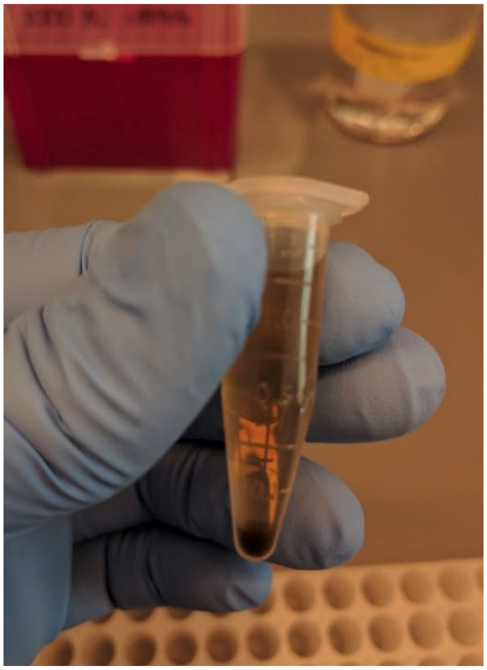

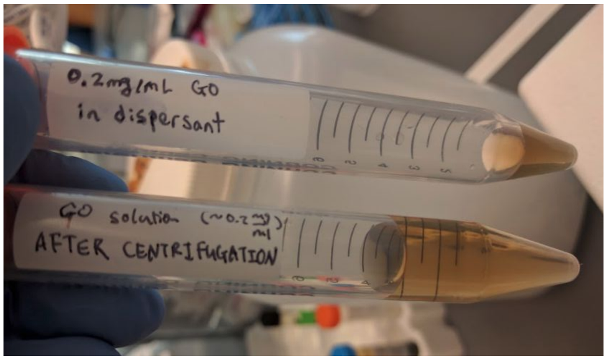


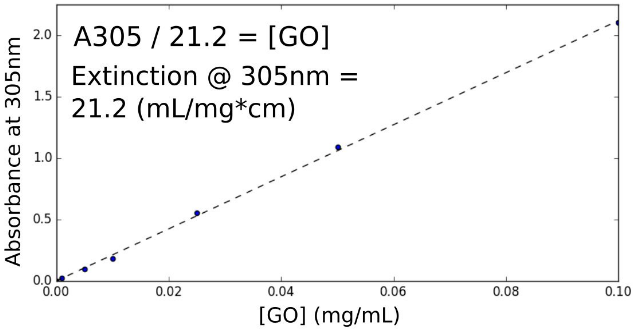


*Preparing the EM grids*

1. Glow discharge Quantifoil 200 mesh 1.2/1.3um EM grids for 30 seconds at 15mA. If a thin plastic film is noticed on the surface of the grids, they should be extensively rinsed in chloroform to remove this film and dried before glow-discharging and coating with GO. If a plasma cleaner is available, we replace glow discharge with a 10 second to a 100% argon plasma.

*Setting up the coating apparatus*

1. Clean a small pyrex petri dish (5 cm diameter) with successive washes of chloroform, ethanol, and milli-Q water. Detergents should be avoided during this wash, as they change the surface tension of the air-water interface.
2. Rinse a copper mesh 'platform' in ethanol and then water. Place the platform into the dish.
3. Add ultrapure water to the dish until the platform is completely covered. This is the subphase. If the platform is too close to the surface, grids will slide when the GO is applied.

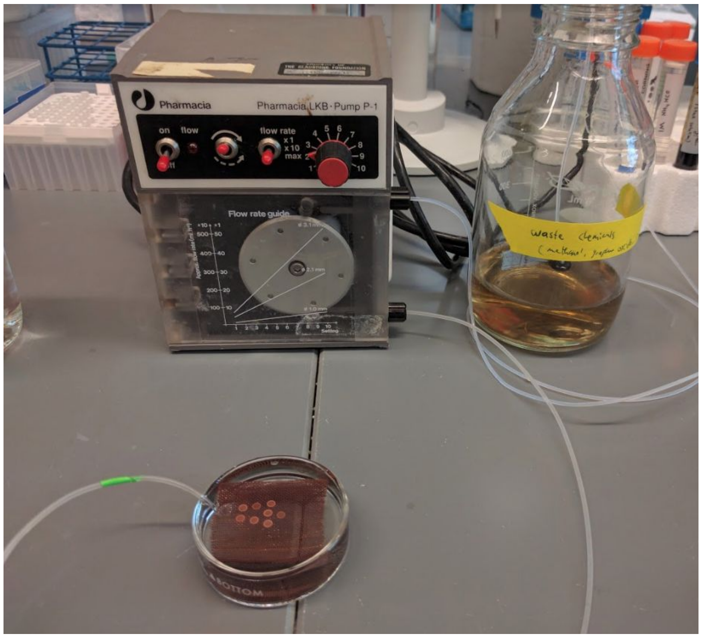

4. Place the inlet line of a peristaltic pump into the dish, under the platform. Place the outlet line into a waste beaker.
5. Carefully submerge grids in the subphase and lie them on the platform, spaced to avoid overlaps. If optimizing the amount of GO applied, one grid is sufficient. Otherwise as many grids can be covered as comfortably fit on the mesh platform.

*Preparing the GO surface and coating the grids*

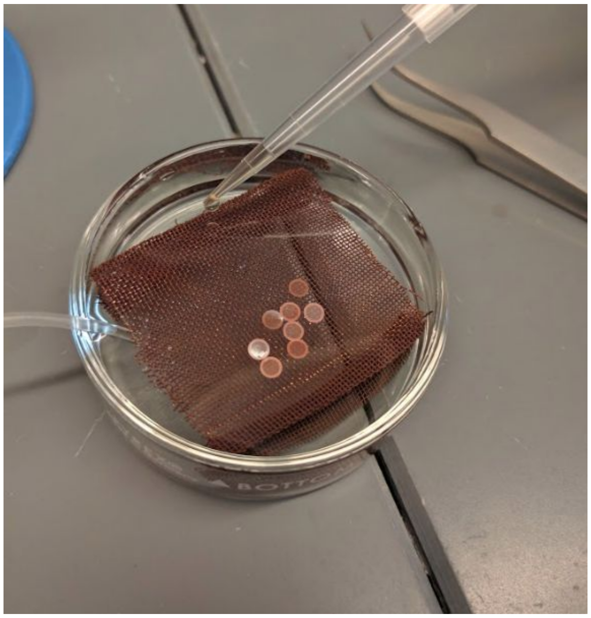

1. Apply the dispersed methanolic GO solution to the surface of the subphase by gently touching successive droplets onto the surface from a pipet or a syringe with a wide gauge needle. The droplet should spread across the surface rather than submerging into the subphase. Apply 10uL droplets. When optimizing the amount of GO, beginning with 400uL (80 micrograms of GO sheets) will likely give overcoated grids, but this can be dialed back in 50uL increments until adequate coating is achieved.
2. With the peristaltic pump, drain the subphase at a flow rate of approximately 1.25 mL/min. The peristaltic pump is used to drain the subphase slowly and steadily: simply aspirating the liquid with a syringe can disrupt the GO film before it has adhered, yielding discontinuous layers.
3. Carefully remove the platform from the petri dish, place it in a larger petri dish, cover and dry overnight. Using grids before they have completely dried will remove the GO layer.
4. Once the grids are completely dry, the GO sheets will be stably adhered to the surface. Any residual subphase can be rinsed before use by applying a droplet of sample buffer and side-blotting on filter paper.

*Storing and cleaning the grids*

Grids can be stored under ambient conditions in a sealed package for several weeks. After several days in air, a layer of contamination can build up on the grids. A short glow discharge (5-10s or less) can usually be tolerated, but caution is advised because longer glow discharges or plasma cleaning will destroy the GO layer.

Evaluating GO coverage in the TEM

1. Add low magnifications (1700-3500x), the size and shapes of GO sheets can be observed.
2. Higher magnifications (~30000x), individual holes can be scrutinized. Optimal holes have few or no high-contrast edges, no aggregates, and complete coverage by the GO film.
3. The number of layers of GO sheets can be determined in diffraction mode. Each GO sheet produces a hexagonal diffraction pattern that may be rotated with respect to other sheets.

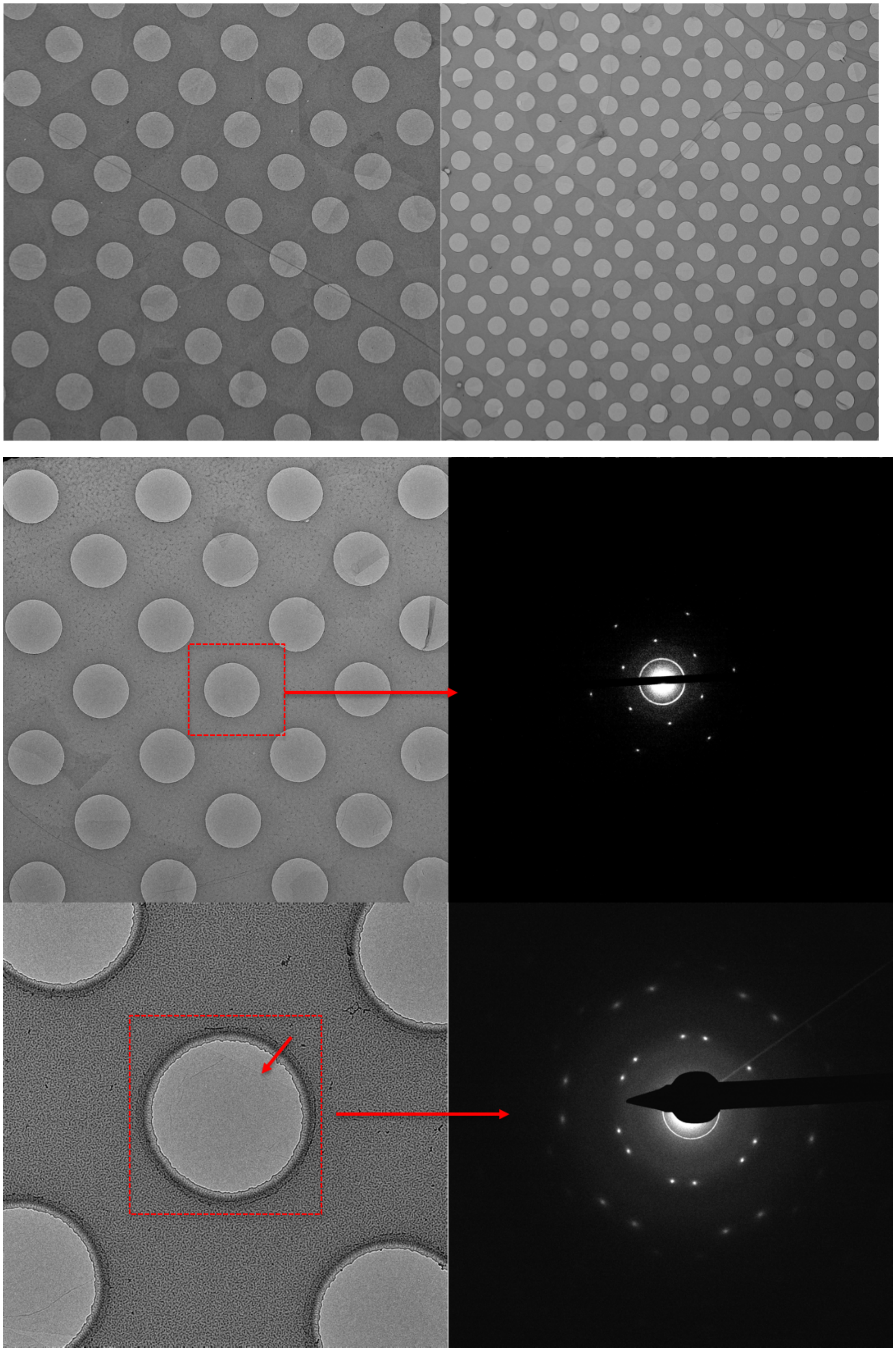
